# CHRONIC LUNG INFLAMMATION LEADS TO MYELOID SKEWING OF HEMATOPOIETIC STEM CELLS IN A CYSTIC FIBROSIS MOUSE MODEL

**DOI:** 10.1101/2024.08.02.606400

**Authors:** Cassia Lisboa Braga, Rubia Isler Mancuso, Evrett N. Thompson, Hasan H. Öz, Ravindra Gudneppanavar, Ping-Xia Zhang, Pamela Huang, Thomas Murray, Marie Egan, Diane S. Krause, Emanuela M. Bruscia

**Affiliations:** Departments of Pediatrics, Yale University School of Medicine, New Haven CT, USA; Laboratory Medicine, Yale University School of Medicine, New Haven CT, USA; Cell Biology, Yale University School of Medicine, New Haven CT, USA; Cellular and Molecular Physiology, Yale University School of Medicine, New Haven CT, USA

## Abstract

Persistent lung inflammation is a hallmark of Cystic fibrosis (CF) lung disease. Inflammation can lead to functional decline in hematopoietic stem cell (HSCs), tipping the balance towards myelopoiesis and contributing to chronic inflammation. However, it’s unknown whether the HSCs are dysfunctional in CF. We tested whether chronic lung inflammation impacts hematopoietic stem and progenitor cells (HSPCs) in a CF mouse model.

Wild-type (WT) and *Cftr*-/- mice were nebulized with lipopolysaccharide (LPS) from Pseudomonas aeruginosa for 5 weeks. The mice were euthanized before the exposure (T0), 24 hours after the last LPS nebulization (T1), or 6 weeks (T2) after the last LPS nebulization. The bone marrow (BM) and lung tissue were collected for flow cytometry analysis of the HSPC population and immune cells in the lungs, respectively. Peripheral blood was collected for complete blood count analysis.

At baseline, *Cftr*-/- mice show a larger HSPC population with a myeloid bias, indicated by higher frequencies of LSK, LT-HSC, CD41^+^ LT-HSC, GMPs, and MkPs. Following chronic LPS nebulization, *Cftr*-/- mice exhibit greater HSPC expansion and myeloid differentiation, alongside increased peripheral myeloid cell counts. Post-recovery, while most HSPC populations return to baseline, *Cftr*-/- mice retain elevated myeloid-biased LT-HSCs, suggesting a persistent myeloid bias. These findings underscore a prominent shift toward myeloid hematopoiesis in CF, which is accentuated by chronic inflammation and remains even after recovery.

Further experiments are underway to assess maladaptive epigenetic changes in HSC as well as if chronic lung inflammation impacts HSC functionality.

## INTRODUCTION

Cystic fibrosis (CF) is a life-threatening genetic disorder that primarily affects the respiratory and digestive systems.^1^ The disease results from mutations in the gene that encodes the cystic fibrosis transmembrane conductance regulator (CFTR), an ATP-gated chloride (Cl-) channel expressed on the apical side of epithelial cells. The lung disease associated with CF is the leading cause of morbidity and mortality in CF patients.^2^ CF lung pathophysiology is hallmarked by a thick and sticky mucus that obstructs airways and makes breathing difficult. This mucus also provides an environment conducive to bacterial growth, leading to chronic lung infections that are difficult to treat.^2^

While CF lung disease is characterized by chronic inflammation and recurrent infections, the role of hematopoiesis in immune cell dysregulation in CF pathophysiology is poorly understood. Hematopoietic stem and progenitor cells (HSPCs) play a critical role in emergency hematopoiesis following the activation of pathogen recognition receptors (PRRs) or via proinflammatory cytokine signals, resulting in the increased production of additional immune cells.^3,4^

The immune system responds to chronic lung infections by producing excessive amounts of inflammatory cells and molecules, which worsens disease outcomes.^5^ In a previous study conducted by our group, Hasan et al. utilized a chronic inflammatory mouse model that mimics the irreversible lung remodeling and functional decline observed in CF patients.^6^ After subjecting *Cftr*-/- mice to chronic nebulization with lipopolysaccharide (LPS) from Pseudomonas aeruginosa for five weeks, followed by a six-week recovery period, they observed significant accumulation of monocytes and neutrophils in the lungs. These immune cells played a role in promoting and perpetuating lung structural damage in *Cftr*-/- mice.^7^ The primary factor driving this lung damage was the excessive recruitment of monocytes, which retained a pro-inflammatory gene signature after six weeks of recovery from chronic LPS. However, given the short lifespan of these cells and the lengthy recovery period, we hypothesize that the inflammatory milieu in CF disrupts hematopoiesis by altering the signals that regulate the differentiation and proliferation of myeloid progenitor cells. Therefore, we analyzed the bone marrow by flow cytometry of WT and *Cftr*-/- mice at baseline (T0), after 5 weeks of chronic LPS exposure (T1), and 6 weeks of recovery following chronic LPS exposure (T2). We demonstrate that the effect of chronic LPS inhalation affects HSPC populations within the bone marrow and has a long-lasting impact on hematopoiesis in CF.

## RESULTS

### *Cftr*-/- mice have a larger population of HSPCs than WT mice, with myeloid dominance at steady state

We used a *Cftr*-/- mouse model^7^ to investigate whether hematopoiesis differs in *Cftr*-/- versus WT mice at baseline (**Figure 1A**,**B**). HSPCs and mature cell populations in the bone marrow were assessed by flow cytometry (**Supplementary Table 1**). Blood was collected for complete blood count (CBC). Within the HSPC pool, the LSK gate contains the hematopoietic stem cells (HSCs) as well as several populations of multipotent progenitors (MPPs) with various lineage biases.^8^ Within the pool of HSCs, long-term HSCs (LT-HSCs) are defined as LSK CD150^+^ CD48^-^ and contain a subpopulation of myeloid-biased LT-HSCs defined as LSK CD150^+^ CD48^-^ CD41^+^ (CD41^+^ LT-HSC).^8^

**Figure 1.**
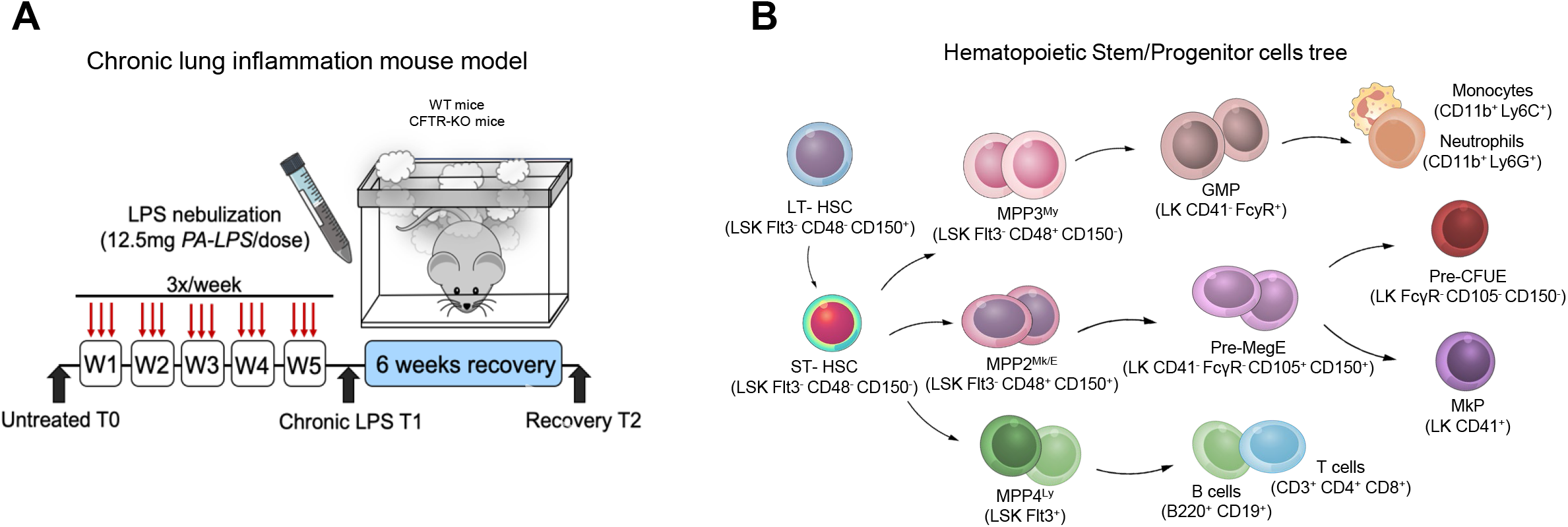
(**A**) Chronic lung inflammation animal model. (**B**) Hematopoietic stem/progenitor cells tree with markers used for flow cytometry analysis.

At baseline, no differences were found in the total number or percentage of lineage negative (Lin^-^) cells (**Figure 2**). However, the LSK, LT-HSC, and CD41^+^ LT-HSC population percentage were higher in *Cftr*-/- than WT mice.

**Figure 2.**
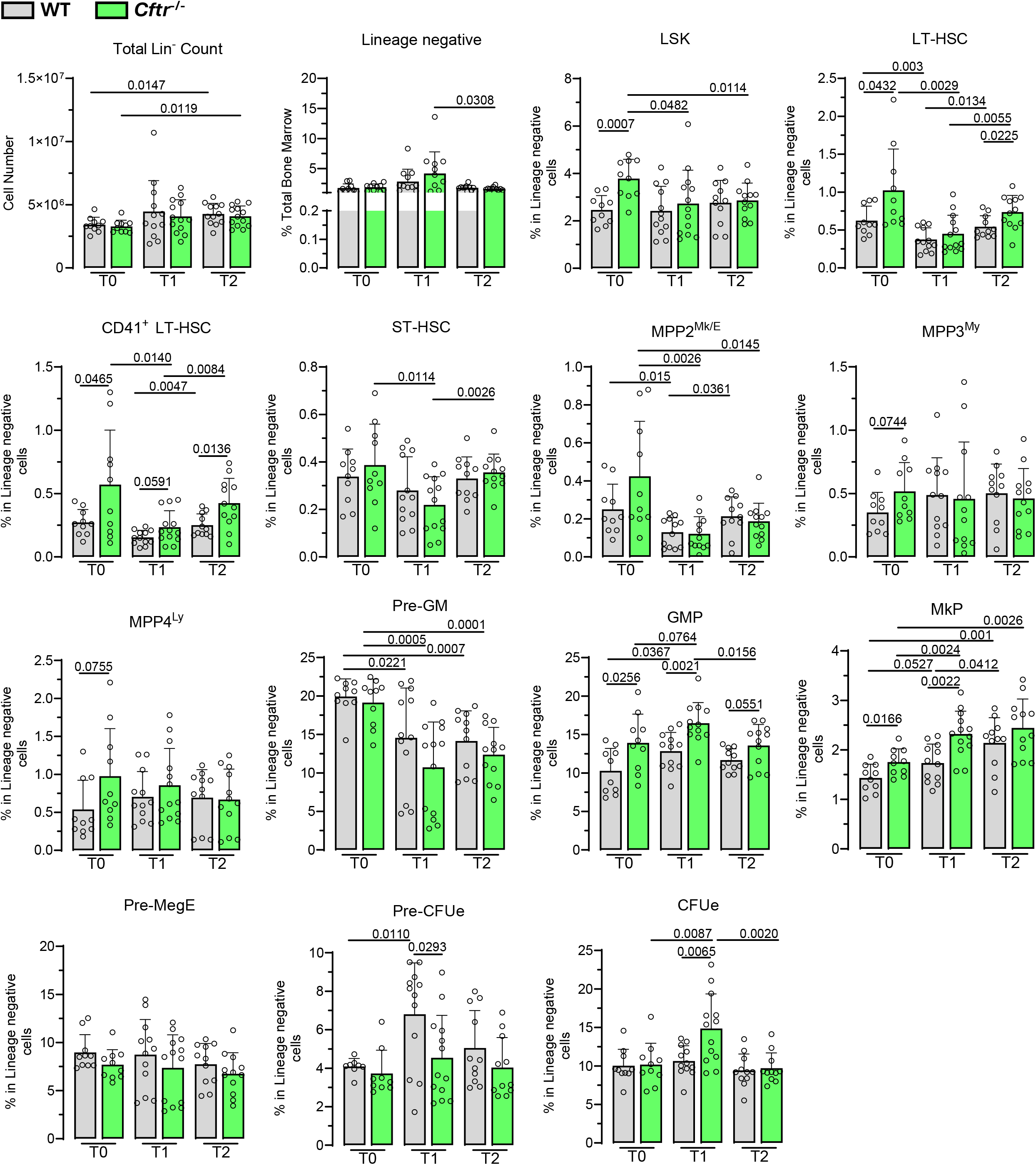
Frequencies of total bone marrow cells and bone marrow progenitors’ populations lineage negative cells, LSK, LT-HSC, CD41^+^ LT-HSC, ST-HSC, MPP2, MPP3, MPP4, Pre-GM, GMP, MkP, Pre-MegE, Pre-CFUe, and CFUe.

No significant differences were observed in the MPPs, although all MPPs showed an increasing trend in *Cftr*-/- compared to WT mice. Analysis of progenitors committed to the myeloid lineage revealed that the percentage of the granulocyte-monocyte progenitor (GMP) and megakaryocyte progenitor (MkP) populations were higher in *Cftr*-/- than in WT (**Figure 2**).

Consistent with the larger frequency of myeloid progenitors in the BM, peripheral blood analysis at baseline (T0) showed that *Cftr*-/- mice had higher monocyte and neutrophil counts compared to WT mice (**Figure 3A**). These data indicate at baseline levels, *Cftr*-/- HSPCs, show a bias towards myeloid cells with a subsequent increase in myeloid output in the blood.

**Figure 3.**
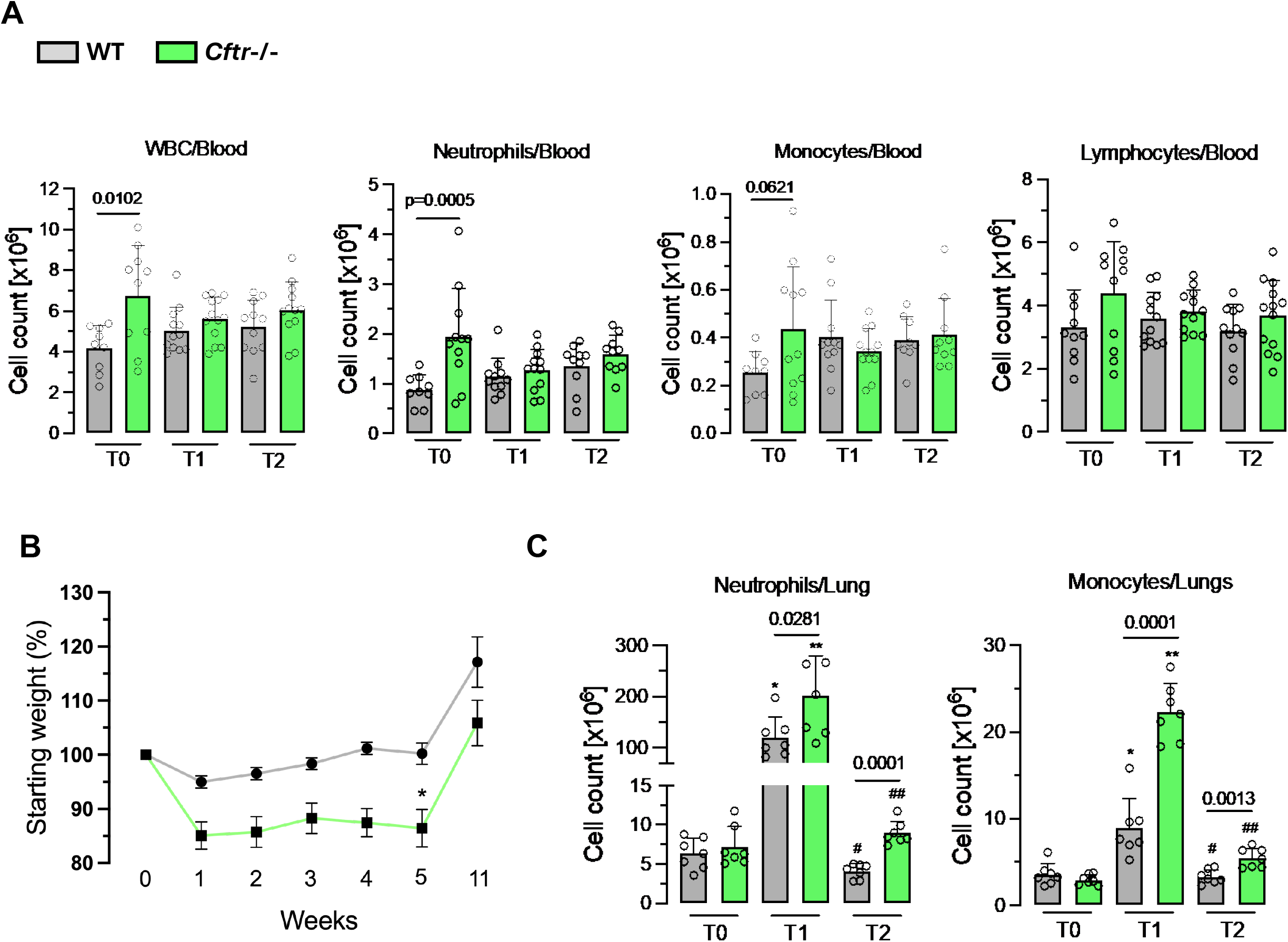
**(A)** White blood cells (WBC), neutrophils, and monocytes count in the peripheral blood, as assessed with Hemavet. **(B)** Weight loss assessed before, during chronic LPS, and 6 weeks after recovery. **(C)** Monocytes and neutrophils count in the lungs assessed by flow cytometry.

### After chronic exposure to LPS, *Cftr*-/- mice exhibit greater HSPC expansion and myeloid differentiation compared to WT mice

To study the effect of chronic lung inflammation on HSPCs, we chronically nebulized WT and *Cftr*-/- mice with LPS (**Figure 1A**). This model mimics the irreversible lung remodeling and decline in lung function observed in individuals with CF.^7^

As previously demonstrated,^6^ *Cftr*-/- mice exhibit a more extensive weight loss and inflammatory response after 5 weeks of nebulization, with higher numbers of monocytes and neutrophils in the lungs (**Figure 3B**).

Immediately after the 5-week LPS challenge (T1), the percentage and total number of Lin^-^ cells in the marrow remains unchanged (**Figure 2**); however, there is a decrease in the frequency of LT-HSC, MPP2, and Pre-GM (**Figure 2**) in both groups when compared to baseline levels. *Cftr*-/- mice specifically demonstrate an additional decrease in LSK, CD41^+^ LT-HSC, and ST-HSC due to the higher baseline levels. The myeloid-biased CD41^+^ LT-HSC subpopulation appears elevated at T1 compared with WT, but this difference does not reach statistical significance. These changes in the percentage of HSPCs align with a high demand for myeloid cell production caused by chronic exposure to LPS. Moreover, it suggests that chronic LPS exposure leads to higher HSPC differentiation in *Cftr*-/- mice. Additionally, the percentage of GMPs and MkPs increases in *Cftr*-/- after the LPS challenge and is higher compared with WT. These observations align with the higher presence of myeloid cells in the lungs of *Cftr*-/- mice compared to WT.

Together, these data suggest that chronic lung inflammation in *Cftr*-/- mice leads to a more prominent HSPC differentiation toward the myeloid lineage.

### *Cftr*-/- mice replenish elevated myeloid-biased LT-HSCs following recovery from chronic LPS exposure

To investigate whether chronic lung inflammation has a long-lasting effect on HSPCs, we analyzed the bone marrow and blood from mice following 6 weeks of recovery after the LPS nebulization (T2). The majority of WT and *Cftr*-/- HSPC populations returned to baseline except for increased total Lin^-^ cell counts, decreased frequency of MPP2, and decreased frequency of Pre-GM. The frequency of MkP were increased specifically in the *Cftr*-/- mice (**Figure 2**). At T2, *Cftr*-/- shows a higher frequency of LT-HSC, CD41^+^ LT-HSC, GMP, and MkP when compared to WT (**Figure 2**).

Interestingly, the myeloid-biased CD41^+^ LT-HSC was higher at T2 in *Cftr*-/- compared to WT, recapitulating the difference in the CD41^+^ LT-HSC population observed at T0. This suggests that even after the reduction of the CD41^+^ LT-HSC to comparable levels between WT and *Cftr*-/- following chronic LPS exposure (T1), an elevated CD41^+^ LT-HSC is a stable characteristic of *Cftr*-/- bone marrow.

In the lungs, despite monocytes’ and neutrophils’ short lifespans, we still observed higher numbers of these cells in *Cftr*-/- compared with WT (**Figure 3C**).

## DISCUSSION

In this study, we used *Cftr*-/- mouse model to study the lung-bone marrow axis and how chronic lung inflammation would impact hematopoiesis. Although CF lung disease is the leading cause of death in people with CF,^9^ the *Cftr*-/- mouse model has limited features of the lung pathology phenotype, potentially because of redundant Cl-channels in the lungs.^10^ To induce a lung phenotype, we developed a mouse model^6,7^ that uses chronic nebulization with LPS to mimic the hyperinflammatory lung environment and lung remodeling observed in patients. The excessive and prolonged recruitment of monocytes and neutrophils, as well as the monocyte gene signature indicating a perpetuation of the immune cells’ chemotaxis, made us wonder if this led to or caused pathological changes in the hematopoietic compartment.

In this study, we found that at the steady state, *Cftr*-/- mice have differences in the hematopoietic system and circulation, that lead to a hyper-response to infection or inflammatory insult. Indeed, the LPS challenge triggered a more significant and prolonged inflammatory response in *Cftr*-/- mice.

The differences observed in HSPC in baseline may result from extrinsic and/or intrinsic factors. BM HSC resides in a specialized BM microenvironment essential for cell activity and regulation.^11^ Once these cells are in intimate physical interaction with surrounding cells like osteoblasts, stromal cells, and vascular cells, malfunctioning of those cells may have an impact on HSPCs.^12^ CF pathology is also characterized by bone disease, marked by a reduction in bone density with reduced osteoclasts and osteoblasts activity.^13^ The CF bone phenotype can also be observed in the *Cftr*-/- mouse model.^14^ Thus, an altered HSC niche, such as the CF bone disease, may impact HSPC independently of infections or inflammatory triggers. Furthermore, the *Cftr*-/- mouse model also recapitulates the features of human CF gastrointestinal disease, such as gut obstruction and inflammation,^10^ which might be an underlying mechanism for altered hematopoiesis at baseline levels.

CFTR expression has also been reported on HSPC.^15^ However, whether CFTR plays a role in hematopoiesis is yet to be determined. Nonetheless, CF patients develop clinically significant anemia, implying a potential role for CFTR in hematopoiesis.^16^

In this *Cftr*-/- mouse model, chronic exposure to LPS exacerbated a pre-existing inflammatory state in *Cftr*-/- mice. This is particularly notable given that chronic LPS exposure led to a significant reduction in certain HSPC populations across both groups, with a more substantial decrease observed in *Cftr*-/- mice. This reduction could reflect an accelerated differentiation of HSPCs into mature myeloid cells in response to chronic inflammation. Indeed, the data show a marked increase in the frequency of myeloid progenitors (GMP and MkP) in *Cftr*-/- mice following LPS exposure in BM and myeloid cells in lungs, reinforcing that CF HSPCs are more responsive to inflammatory signals.

HSPCs can quickly adjust to inflammatory insults to meet increased demand.^17^ While this adaptive response may be advantageous in promoting resistance to infection, it also has the potential to contribute to the perpetuation of chronic inflammatory conditions through the establishment of a deleterious feed-forward mechanism.^18^ Notably, studies indicate that chronic inflammation may have a long-lasting impact on HSCs by facilitating the acquisition of epigenetic maladaptive memory, thereby leading to compromised HSC functionality and eventual exhaustion of these cells.^19^

Since persistent infections and chronic lung inflammation are hallmarks of CF disease, we speculate that the chronic aspect of the lung disease may be, at least, partly fueled by changes affecting HSCs, which sustain inflammation and immune dysregulation in CF. Previous reports have shown that systemic LPS can activate LT-HSCs.^20^ Here, we provide evidence that chronic lung inflammation in *Cftr*-/-, induced by prolonged LPS exposure, affects HSPCs in the BM.

Following a six-week recovery period after LPS exposure, *Cftr*-/- mice retained a higher frequency of LT-HSCs, CD41^+^ LT-HSCs, GMPs, and myeloid cells in the lungs than WT mice. Despite the overall normalization of HSPC numbers, the sustained elevation in these populations suggests a form of epigenetic memory or long-term alteration in the hematopoietic system. These alterations may predispose *Cftr*-/- mice to a heightened inflammatory response upon subsequent insults, potentially exacerbating disease progression. However, whether chronic lung inflammation promoted maladaptive innate trained immunity on HSC is yet to be determined.

In conclusion, our study reveals that *Cftr*-/- mice have altered hematopoiesis at steady-state, and chronic lung inflammation led to an exacerbation of a previous pro-inflammatory state. The observed dysregulation in hematopoiesis and the retention of myeloid cells in *Cftr*-/- mice imply that chronic lung inflammation has far-reaching effects beyond the immediate inflammatory response. These alterations in HSPCs could contribute to the chronic inflammation observed in CF by continuously supplying pro-inflammatory myeloid cells to the circulation. This could create a feedback loop that maintains or exacerbates lung inflammation, thus worsening disease outcomes.

These findings underscore the importance of considering hematopoietic changes in the broader context of CF pathophysiology.

## METHODS

### Mouse models and in vivo studies

All procedures followed the relevant laws and institutional guidelines and were approved by the Yale University Institutional Animal Care and Use Committee. The transgenic *Cftr*-/- (B6.129P2-KOCftrtm1UNC) mice were obtained from The Jackson Laboratory and bred in the Yale University Animal Facility. *Cftr*-/- mice were given drinking water supplemented with Colyte (Braintree Labs) to prevent gastrointestinal obstruction at weaning. Non-CF littermate controls were kept on the same dietary conditions as the CF mice to avoid nutritional confounding. All mice were 4 to 6 months old at the beginning of the experiment, and an equal number of female and male mice were included in the study.

### Chronic lung inflammation mouse model

*Cftr*-/- mice and *Cftr*+/+ (WT) were subjected to chronic nebulization using a Pulmo-Aide Compressor (Natallergy, Duluth, GA) with lipopolysaccharide (LPS) from Pseudomonas aeruginosa (L9143 from Millipore Sigma). The mice received 12.5 mg of LPS per 5ml of PBS for 15 minutes, 3 times a week, over five weeks. The mice were euthanized by intraperitoneal injection of urethane 8% in PBS before the chronic LPS exposure (T0), 24 hours after the last nebulization (T1), or after 6 weeks of recovery from LPS (T2) (**Figure 1A**).

### Peripheral blood collection

At times T0, T1 and, T2, up to 500 μl of peripheral blood was collected from the orbital vein. A complete blood count (CBC) was assessed using automate blood count (Hemavet 950 Drew Scientific) with 50 μl of blood.

### Lung collection and preparation for flow cytometry

The lungs were collected by making a midline incision from the sternum to the diaphragm, and blood was removed from the pulmonary circulation by perfusing with PBS supplemented with heparin (1:1000) through the heart. The right inferior lobes were then collected in PBS and used for flow cytometry.

The lung samples were processed using a dissociation kit (Miltenyi, CA) and a GentleMACS dissociator (Miltenyi, CA) following the manufacturer’s instructions, to obtain single cell suspensions. Afterward, 1×10^6^ cells were stained with a fixable viability dye (Invitrogen, MA), incubated with FcBlock (BD Biosciences, CA), and then washed and stained with freshly prepared antibody cocktails for 30 minutes at 4°C, in accordance with standard cell surface staining protocols.

After washing and fixing the cells with BD Cytofix (BD Biosciences, CA), samples were analyzed using a BD LSR II with BD FACSDiva software. Data analysis was carried out using FlowJo software (TreeStar, Ashland, OR). To obtain the total live cell counts, the percentage of each assessed population among singlet/live cells was multiplied by the total lung cell counts in the inferior lobes.

### Bone marrow cell isolation and lineage depletion

Femur, tibia, pelvic and spine bones were isolated from euthanized mice and crushed with a pestle. The cell suspension was filtered through 100 mm cell strainer (Greiner Bio-one) using FACS buffer. The whole bone marrow was depleted using the EasySep™ Mouse Hematopoietic Progenitors Cell Isolation Kit (#19856) according to the manufacturer’s instructions. Samples were stained with a cocktail of antibodies and analyzed using a BD FACSAria with BD FACSDiva software (**Supplementary Table 2**).

