## Supplemental Table 1 for "CHRONIC LUNG INFLAMMATION LEADS TO MYELOID SKEWING OF HEMATOPOIETIC STEM CELLS IN A CYSTIC FIBROSIS MOUSE MODEL"

|  | **Cell Population** | **Flow Cytometry Markers** |
| --- | --- | --- |
| LT-HSC | Long-term Hematopoietic Stem Cells | LSK Flt3^-^ CD48^-^ CD150^+^ |
| CD41^+^LT-HSC | Long-term Hematopoietic Stem Cells CD41^+^ | LSK Flt3^-^ CD48^-^ CD150^+^ CD41^+^ |
| ST-HSC | Short-term Hematopoietic Stem Cells | LSK Flt3^-^ CD48^-^ CD150^-^ |
| MPP2 | Multipotent Progenitors 2 | LSK Flt3^-^ CD48^+^ CD150^+^ |
| MPP3 | Multipotent Progenitors 3 | LSK Flt3^-^ CD48^+^ CD150^-^ |
| MPP4 | Multipotent Progenitors 4 | LSK Flt3^+^ |
| Pre-GM | Pre-Granulocytes/Macrophages Progenitors | Lin^-^ Sca1^-^ Kit^+^ CD41^-^ FcγR^-^ CD105^-^ CD150^-^ |
| GMP | Granulocytes/Macrophages Progenitors | Lin^-^ Sca1^-^ Kit^+^ CD41^-^ FcγR^+^ |
| Pre-MegE | Pre-Megakaryocyte/Erythroid Progenitors | Lin^-^ Sca1^-^ Kit^+^ CD41^-^ FcγR^-^ CD105^-^ CD150^+^ |
| Pre-CFUE | Pre-Colony Forming Unit-Erythroid | Lin^-^ Sca1^-^ Kit^+^ CD41^-^ FcγR^-^ CD105^+^ CD150^+^ |
| CFUE | Colony Forming Unit-Erythroid | Lin^-^ Sca1^-^ Kit^+^ CD41^-^ FcγR^-^ CD105^+^ CD150^-^ |
| MkP | Megakaryocytic Progenitors | Lin^-^ Sca1^-^ Kit^+^ CD150^+^ CD41^+^ |
