## Supplemental Table 2 for "CHRONIC LUNG INFLAMMATION LEADS TO MYELOID SKEWING OF HEMATOPOIETIC STEM CELLS IN A CYSTIC FIBROSIS MOUSE MODEL"

**Supplementary Table 2**. Flow cytometry antibodies used for HSPC panel.

| **Antibody** | **Fluorophore** | **Clone** | **Company** | **Catalog #** |
| --- | --- | --- | --- | --- |
| Anti-mouse CD3ε | PE/Cyanine7 | 145-2C11 | Biolegend | 100320 |
| Anti-mouse CD11b | PE/Cyanine7 | M1/70 | Biolegend | 101216 |
| Anti-mouse CD45R/B220 | PE/Cyanine7 | RA3-6B2 | Biolegend | 103222 |
| Anti-mouse Ly-6G/Ly-6C (Gr-1) | PE/Cyanine7 | RB6-8C5 | Biolegend | 108416 |
| Anti-mouse TER-119/Erythroid cells | PE/Cyanine7 | TER-119 | Biolegend | 116222 |
| Anti-mouse CD117 | APC | 2B8 | Biolegend | 105812 |
| Anti-mouse Ly-6A/E (Sca-1) | PerCP | D7 | Biolegend | 108122 |
| Anti-mouse CD135 (FLT3) | PE | A2F10.1 | BD Biosciences | BD553842 |
| Anti-mouse CD48 | APC-Cyanine7 | HM48-1 | Biolegend | 103432 |
| Anti-mouse CD150 (SLAM) | Alexa Fluor 488 | TC15-12F12.2 | Biolegend | 115916 |
| Anti-mouse CD41 | PE/Dazzle™ 594 | MWReg30 | Biolegend | 133936 |
| Anti-mouse CD16/CD32 | BUV737 | 2.4G2 | BD Biosciences | BD612783 |
| Anti-mouse CD105 | Brilliant Violet 421 | MJ7/18 | BD Biosciences | 562760 |
| Anti-mouse CD127 (Il-7Rα) | Brilliant Violet 605 | A7R34 | Biolegend | 135041 |
